# Brain Mechanisms of Lexicality in Bilingual Emotion Processing

**DOI:** 10.64898/2026.09.21.753217

**Authors:** Cheng Xiao, Jiang Liu, Yuqi Zhang, Wanzhi Lyu, Rutvik H. Desai, Ma Feilong

**Affiliations:** Department of Psychology, University of South Carolina; Linguistics Program, University of South Carolina; Department of Language, Literatures and Cultures, University of South Carolina; Department of Psychological and Brain Sciences, Dartmouth College; Institute for Mind and Brain, University of South Carolina

## Abstract

Understanding how native speakers and second language (L2) learners process language differently is fundamental to theories of language acquisition, comprehension, and use. A central debate in this literature concerns the inconsistent findings in emotional prosody processing between native speakers and L2 learners, partly attributable to stimulus lexicality: some studies use real words, whereas others use pseudowords. To resolve this discrepancy, we measured fMRI responses in 15 native speakers and 15 L2 learners while they listened to Mandarin real-word and pseudoword stimuli. Both groups engaged in overlapping auditory, cognitive, and affective regions, including the auditory cortex, inferior frontal cortex, superior temporal cortex, and the temporoparietal junction. However, the two groups showed distinct responses depending on lexicality: native speakers showed stronger recruitment of temporal regions for pseudowords, whereas L2 learners showed greater frontoparietal activation for real words. These patterns were consistent across positive and negative prosody conditions. Furthermore, among L2 learners, higher proficiency was associated with greater frontal activation for real words, but not pseudowords. Together, these findings demonstrate that the differences between native speakers and L2 learners depend critically on stimulus lexicality, highlighting the role of lexicality in bilingual emotion processing.

## Introduction

Imagine hearing a newly coined slang term for the first time. Even before you know what it means, you can often tell that it sounds word-like yet unfamiliar. This everyday experience highlights a fundamental distinction in language processing: some spoken forms readily activate existing lexical and semantic representations, whereas unfamiliar word-like forms do not. This distinction may shape not only how we recognize words, but also how we extract other information carried by speech, such as a speaker’s emotional tone. How does the brain process emotional speech when lexical meaning is available versus when it is not, and does this distinction matter more for some listeners than others?

Emotional information in speech is conveyed not merely through what is said, but also through how it is said ^1–4^. This “how” can be carried by emotional prosody, the modulation of tone of voice to express feelings and attitudes ^5^, which links linguistic expression to emotional experience across a range of social interactions ^6–8^. Successful communication requires listeners to integrate linguistic and paralinguistic information simultaneously within milliseconds, and this integration becomes especially demanding in a second language (L2) ^9–11^.

A substantial body of literature has established that L2 learners are often less efficient than native speakers in both emotion ^3,12–16^ and word processing ^17–22^. L2 words are consistently recognized less efficiently than native words, attributed to either weaker lexical–conceptual links ^23^ or competition between lexical candidates across native and second languages ^24^. In emotion processing, however, the picture is less clear. The In-Group Advantage (IGA) hypothesis ^3,16^ posits that although emotional expressions, including emotional prosody, can often be recognized across cultural groups at above-chance levels, native speakers have an advantage in recognizing emotional prosody when expressed by members of one’s own cultural group. However, empirical evidence for the IGA effect remains mixed: some studies show that native listeners outperform non-native or L2 listeners ^15,25–27^, whereas others find comparable performance ^28,29^, and some even report L2 advantages for particular emotion types ^30,31^.

One source of these mixed results is the lexicality of the stimuli, namely, the presence or absence of lexical-semantic representation (real words vs. pseudowords). Some studies use real words and real-word sentences, requiring listeners to process lexico-semantic and emotional prosody concurrently ^30,31^, while others use pseudowords which removes lexico-semantic content and isolates emotional prosody processing on its own ^27,32^. This lexicality effect is further confounded by listeners’ semantic knowledge of the stimuli ^33,34^: native speakers can reliably access the meaning of most stimuli, whereas L2 learners’ semantic knowledge is often incomplete or variable, suggesting that observed group differences may also reflect unequal word recognition rather than differences in emotional prosody processing. Therefore, we controlled for the semantic knowledge of native speakers and L2 learners to examine emotional prosody processing in both real words and pseudowords.

Neuroimaging research has revealed similarly mixed findings. For emotional prosody, while classic models primarily attributed emotional prosody to the right hemisphere ^35–37^, contemporary meta-analyses instead pointed to a bilateral network ^4,7,38,39^ involving superior temporal cortex, auditory, and frontal regions. For lexicality, some studies found no differences in real word and pseudoword processing ^40,41^, while others reported differences in temporal and inferior parietal regions ^42–46^. Despite this, emerging evidence shows that lexicality and emotional prosody interact. Behaviorally, native speakers recognized emotional prosody faster in pseudoword than real-word sentences ^47^. Such lexicality difference is further modulated by the congruence between semantics and emotional prosody ^15,48^. Electrophysiologically, emotional prosody-only expectancy violations in pseudoword sentences elicited a right-lateralized positive-going response, whereas violations involving the integration of emotional prosody with lexicality in real-word sentences produced a broadly distributed negative N400 response ^49,50^. Additionally, the emotional prosody processing network in real-word sentences mirrors sentence comprehension, with additional bilateral regions engaged when semantic content had to be integrated ^1^. However, few studies have directly tested how lexicality interacts with emotional prosody from a cross-linguistic perspective. It thus remains unknown how lexicality influences emotional prosody processing in the brain across native speakers and L2 learners.

The effect of lexicality on emotional prosody processing is particularly crucial for tonal languages, where the same acoustic feature, pitch, encodes both emotional prosody and lexical meaning ^51^. In Mandarin Chinese, four lexical tones differentiate word meanings: for example, the syllable “ma” means *mother* in Tone 1, *hemp* in Tone 2, *horse* in Tone 3, and *to scold* in Tone 4 ^52^. In tonal languages, the real-word and pseudoword distinction therefore extends beyond semantics: lexicality determines not only what meaning is available but also how the acoustic signal itself is interpreted. Existing literature reliably shows group differences between native and non-native Chinese speakers in emotion and lexicality processing. For example, native Chinese speakers recruited more left-lateralized frontoparietal activation than native English speakers when processing semantically neutral real-word sentences ^53^. Within L2 Chinese learners, proficiency further shapes these patterns: more proficient learners showed greater left posterior superior temporal gyrus (STG) activation than less proficient learners ^54^. However, none of these studies examined emotional prosody, leaving the neural interplay between lexicality and emotional prosody in tonal languages untested.

Therefore, the present study uses functional magnetic resonance imaging (fMRI) to investigate how native speakers and L2 learners process emotional prosody in Mandarin real words and pseudowords. Pseudowords were constructed using lexical gaps, with matched psycholinguistic properties to real words, and the two groups had similar semantic knowledge for both real words and pseudowords. Fifteen native Chinese speakers and 15 L1-English L2-Chinese learners listened to Mandarin real-word and pseudoword stimuli spoken with positive or negative emotional prosody while undergoing fMRI across four functional runs lasting approximately 45 minutes in total, and performed an emotional prosody judgment task. fMRI data were preprocessed using fMRIPrep ^55^ and functionally aligned across participants using hyperalignment ^56,57^. Specifically, we compared neural responses to real words and pseudowords between native speakers and L2 learners, examined positive and negative prosody separately to test whether lexicality effects were specific to a particular emotion type, and categorized L2 learners into higher and lower proficiency subgroups to examine whether L2 proficiency modulates this lexicality effect. We conducted both whole-brain and ROI analyses to clarify how lexicality shapes the neural differences between native speakers and L2 learners in emotional prosody processing.

## Results IGA effects

To examine the effect of In-Group Advantage (IGA) in recognizing Chinese emotional prosody, native Chinese speakers and L2 Chinese learners completed an emotion judgment task using both real-word and pseudoword stimuli. Each participant completed four 11-minute functional runs, with each run comprising 24 randomized 25.6-second blocks. Within each block, participants were asked to listen to 12 auditory stimuli and then judge whether the speaker conveyed negative or positive emotions. As shown in Figure 1, native and learner groups exhibited above-chance accuracy (ACC) in recognizing emotional prosody for both real-word (*M* = 76.7%, *SD* = 0.423) and pseudoword stimuli (*M* = 78.5%, *SD* = 0.411). Both groups also recognized the emotional prosody rapidly, with mean reaction times (RT) of 597 mgs for real-word stimuli and 623 ms for pseudoword stimuli. We used logistic and linear mixed-effects models to analyze ACC and response time RT, respectively, with participant included as a random intercept. In terms of ACC, native speakers recognized emotional prosody significantly more accurately than L2 learners for both real-word stimuli (*B* = 1.671, *SE* = 0.555, *z* = 3.009, *p* = .003) and pseudoword stimuli (*B* = 1.321, *SE* = 0.495, *z* = 2.670, *p* = .008), supporting the IGA hypothesis in this tonal language. In terms of RT, while no IGA effect was observed, there was a significant lexicality effect: real-word stimuli were recognized significantly faster than pseudoword stimuli (*B* = -31.33, *SE* = 14.52, *t* = -2.158, *p* = .031). No significant interaction between group and lexicality emerged for either ACC or RT. Taken together, the IGA effect appeared in ACC while the lexicality effect appeared in RT, suggesting that group and lexicality may both modulate emotional prosody processing. We therefore examined these effects in the whole-brain analyses below.

**Figure 1.**
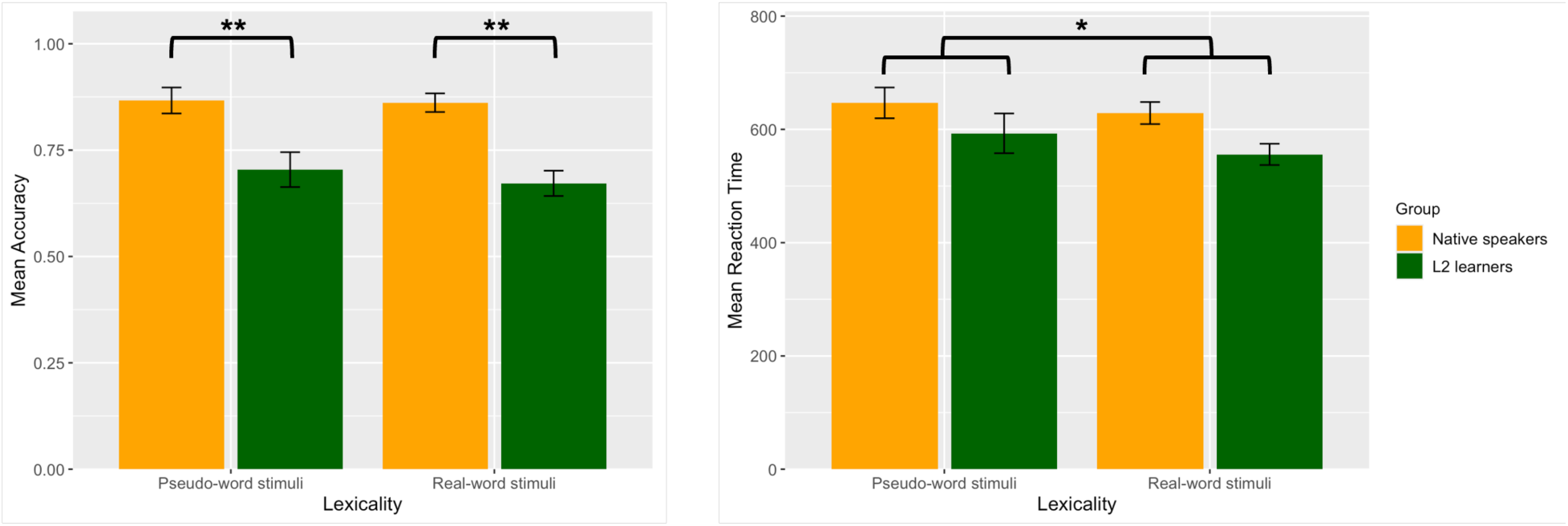
Mean accuracy (left) and reaction time (right) of emotional prosody judgments in the pseudoword and real-word stimuli. Error bars represent 95% confidence intervals. ** *p <* .01, * *p <* .05.

## Interaction effects between lexicality and group

We conducted whole-brain analyses testing the lexicality × group interaction to investigate how lexicality modulates emotional prosody processing in native speakers and L2 learners. We found a significant interaction effect such that native speakers and L2 learners showed distinct brain response patterns to lexicality (*t* = 6.041, *p* < .001). As shown in Figure 2, when processing real words, compared to L2 learners, native speakers showed increased activation in the angular gyrus (AG), a region implicated in lexico-semantic integration ^58,59^, and decreased activation in the middle frontal gyrus (MFG), a region associated with multiple demand processing ^60^. When processing pseudowords, native speakers showed decreased activation in the AG and increased activation in the MFG compared to L2 learners. This pattern suggests that native speakers recruit semantic integration when lexical content is available but shift toward more effortful, top-down processing when it is not, whereas learners show the reverse pattern.

**Figure 2.**
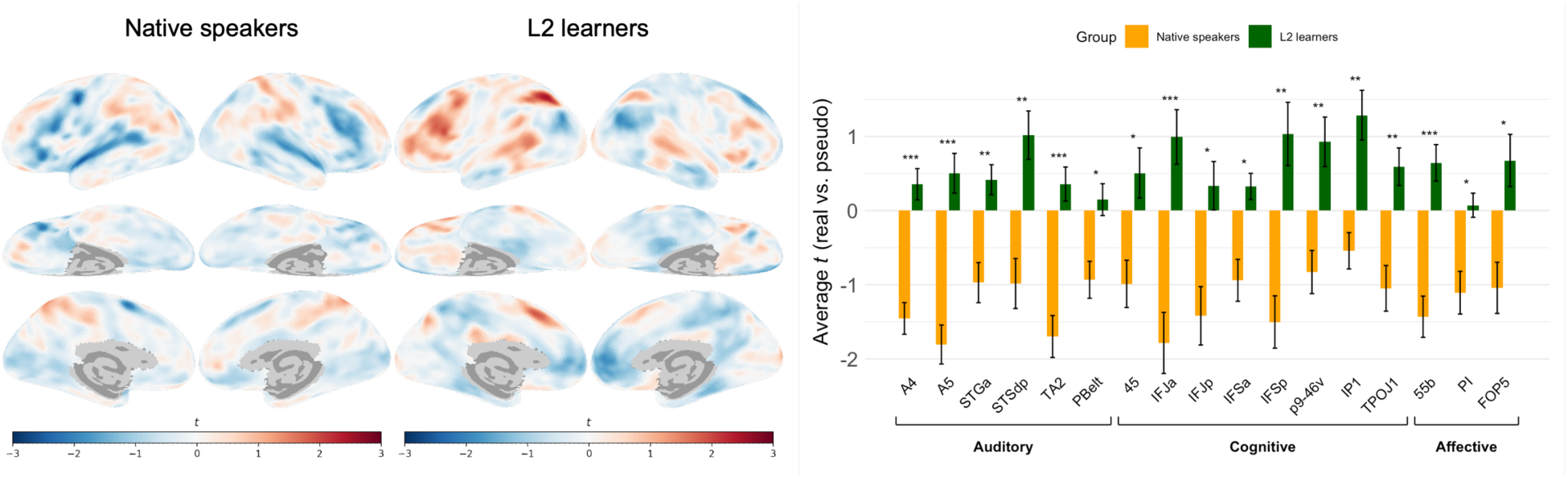
Native speakers and L2 learners show distinct brain response patterns to lexicality. (Left) Contrast maps between real words and pseudowords for native speakers and L2 learners, where the red color indicates greater responses to real words than pseudowords, and blue indicates greater responses to pseudowords. (Right) Average contrast between real words and pseudowords in auditory, cognitive, and affective ROIs for both native speakers (orange) and L2 learners (green). Error bars represent standard errors. \*\*\**p* < .001, \*\**p* < .01, \**p* < .05.

ROI analyses further identified 17 parcels in the real-word vs. pseudoword contrast, clustering in *auditory* regions (A4, A5, STSdp, STGa, TA2, PBelt); *cognitive* regions, including inferior frontal regions (45, IFJa, IFJp, IFSa, IFSp), dorsolateral prefrontal cortex (p9-46v), and inferior parietal region and temporo-parieto-occipital junction (IP1, TPOJ1); *affective* regions, including a music perception hub 55b ^61^, and posterior insula cortex (PI) and frontal opercular area 5 (FOP5). Here, the term “affective” is used narrowly to denote regions implicated in prosodic and salience-related affective processing. This distributed auditory, cognitive, and affective network is broadly consistent with the three-stage model of emotional prosody processing in non-tonal languages ^4^. Additionally, across all parcels, native speakers showed greater engagement for pseudowords, whereas learners showed greater engagement for real words, which parallels the AG/MFG pattern observed at the whole-brain level, further supporting that lexicality differentially modulates the neural processing of emotional prosody in native speakers and L2 learners.

To assess the robustness of these effects, we repeated the analyses in Figure 2 separately for positive prosody and negative prosody. Specifically, we examined the group × lexicality interaction within each prosodic condition (Figure 3). The real-word vs. pseudoword contrast showed largely aligned spatial distributions and effect directions across both prosodic conditions. ROI analyses showed significant group differences in six parcels in auditory and cognitive clusters (A4, A5, STSdp, TA2, IFJa, and IFSp) that operated independently of prosodic valence.

**Figure 3.**
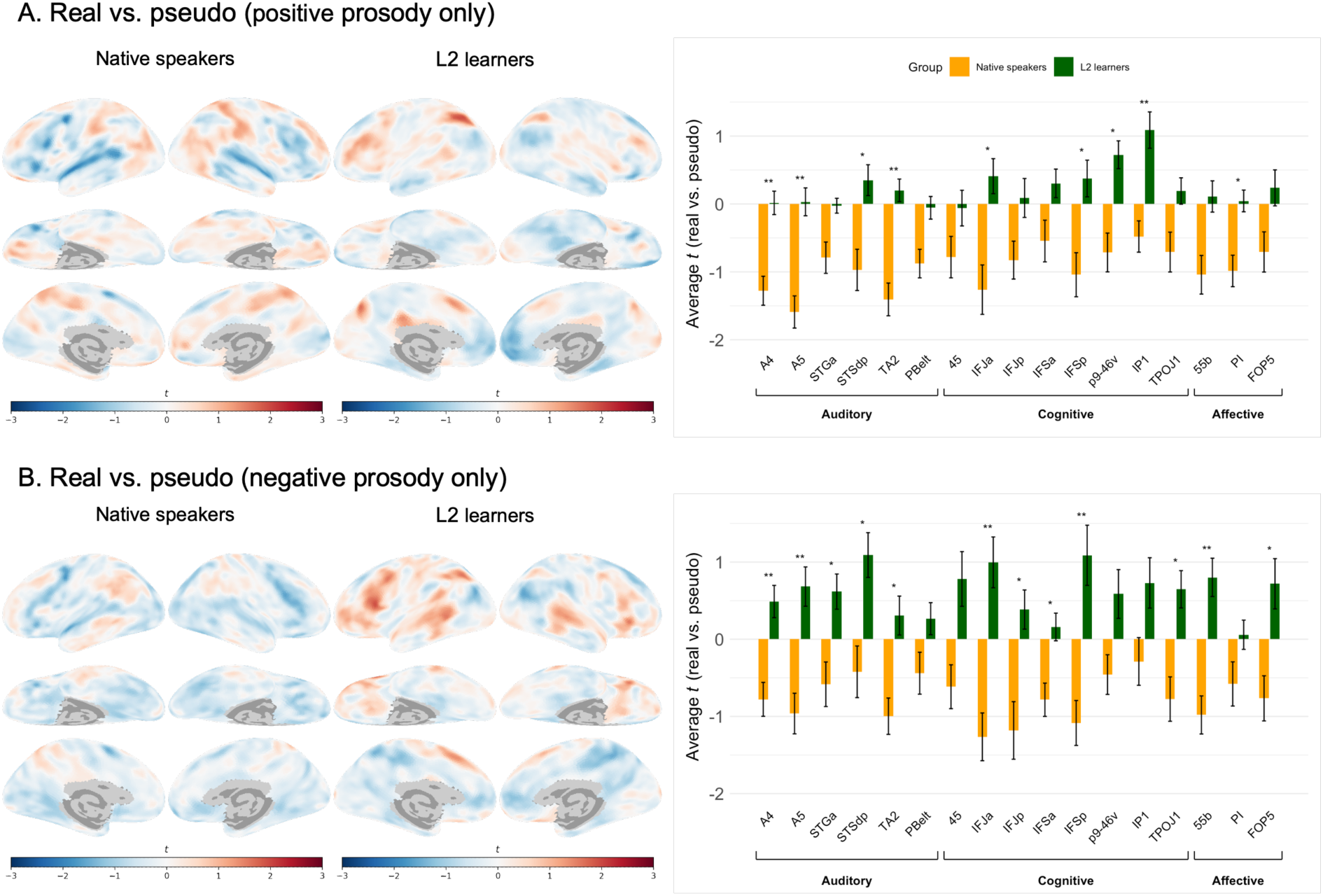
Native speakers and L2 learners show distinct brain response patterns to lexicality for both positive prosody and negative prosody. We repeated the analyses in Figure 2 with only positive prosody and negative prosody and obtained consistent results. (Left) Contrast maps between real words and pseudowords in A) positive prosody only and B) negative prosody only for native speakers and L2 learners, where the red color indicates greater responses to real words than pseudowords, and blue indicates greater responses to pseudowords. (Right) Average contrast between real words and pseudowords in A) positive prosody only and B) negative prosody only in auditory, cognitive, and affective ROIs for both native speakers (orange) and L2 learners (green). Error bars represent standard errors. \*\*\**p* < .001, \*\**p* < .01, \**p* < .05.

In these temporal-frontal regions, the interaction was driven by a greater real-word over pseudoword response in L2 learners compared to native speakers, suggesting a higher cognitive demand for successful lexical access in a second language. Beyond this shared network, we observed valence-specific recruitment: positive prosody uniquely recruited frontoparietal regions associated with cognitive and affective clusters (p9-46v, IP1, and PI), while negative prosody additionally elicited broader engagement across the auditory-cognitive-affective network (STGa, IFJp, IFSa, TPOJ1, 55b, and FOP5). Taken together, these results indicate that processing emotional prosody in real words vs. pseudowords relies on a stable auditory–cognitive–affective network identified in Figure 2.

## Lexicality effects

To clarify the neural basis of the lexicality effect, we compared brain activation between native speakers and L2 learners during their processing of emotional prosody in real words and pseudowords. When processing emotional prosody in real words (Figure 4A), both groups showed widespread activations in frontal, temporal, and parietal regions relative to resting baseline, consistent with engagement of the auditory–cognitive–affective network. ROI results indicated that L2 learners showed stronger activation than native speakers across a set of cognitive and affective regions, including TPOJ1, IFJa, IFJp, IFSa, IFSp, p9-46v, and 55b. L2 learners also showed stronger activation in an additional set of attention and salience regions, including PEF (premotor eye field), AVI (anterior ventral insula), LIPd (lateral intraparietal, dorsal), IP2 (intraparietal area 2), and 8BM (dorsomedial prefrontal). None of the significant regions fell within the auditory network, suggesting that the two groups did not differ in the auditory processing of real words. Notably, in PGi (inferior parietal lobule), native speakers showed stronger activation than L2 learners, suggesting stronger semantic association and integration. These findings reveal that native speakers may process real words more efficiently through semantic-integration mechanisms, whereas L2 learners need to recruit additional cognitive, affective, and attentional resources to accomplish the same task.

**Figure 4.**
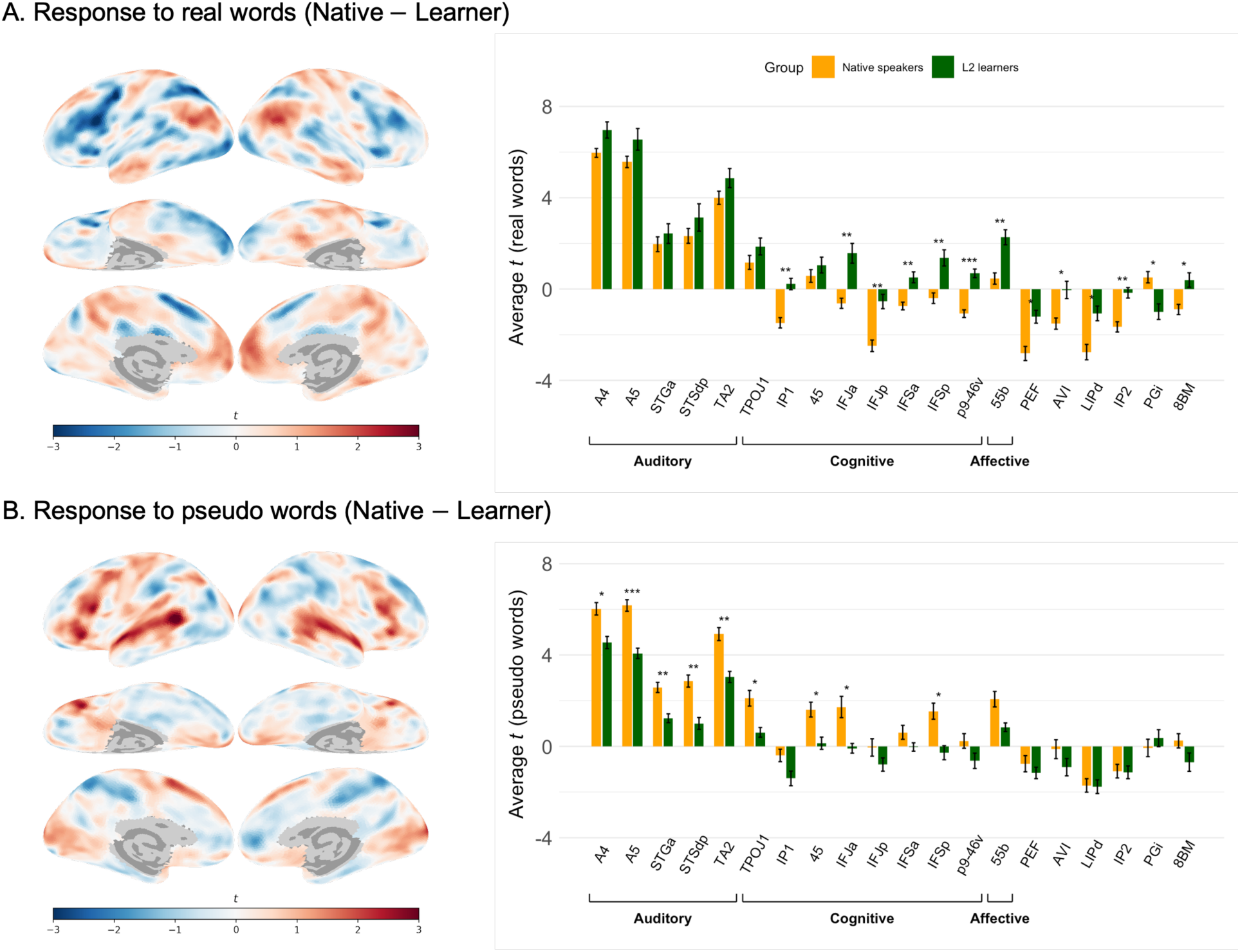
Differences between native speakers and L2 learners when processing real words (top) and pseudowords (bottom). (Left) Contrast maps for A) real words and B) pseudowords showing regions where native speakers exhibit greater activation than L2 learners (red) and where L2 learners exhibit greater activation than native speakers (blue), each relative to resting baseline. (Right) Average contrast in auditory, cognitive, and affective ROIs for both native speakers (orange) and L2 learners (green): A) between real words and resting baseline and B) average contrast between pseudowords and resting baseline. Error bars represent standard errors. \*\*\**p* < .001, \*\**p* < .01, \**p* < .05.

ROI analyses identified three parcels, TPOJ1, IFJa, and IFSa, that were significant for both real words and pseudowords. In all three regions, native speakers showed stronger activation than L2 learners, reversing the direction observed for real words. In addition, as shown in Figure 4B, auditory regions showed significant native > L2 differences, including primary/belt auditory cortex (A4, A5, TA2) and superior temporal regions (STGa, STSdp). Area 45 also showed a significant group difference (native > L2) in processing pseudowords. Because our pseudoword stimuli were phonotactically word-like, this pattern suggests that native speakers may engage semantic selection among phonologically similar real-word neighbors. Accordingly, this process may be less accessible to L2 learners, given their sparser lexical networks.

Together, the observed asymmetry in real words and pseudowords revealed that in the absence of lexical content, native speakers engaged auditory networks more effectively when processing unfamiliar sound sequences, whereas L2 learners showed no comparable engagement of these networks. This distinction is critical because prior work has often reported mixed findings when evaluating native speakers’ advantage in emotional prosody recognition. Our findings suggest that this IGA effect may not be uniform but instead depends on lexicality.

## Lexicality effects and language proficiency

To examine the proficiency effect, we analyzed whether the lexicality effect differed between higher and lower proficiency learners. We categorized L2 learners into higher and lower proficiency subgroups based on their averaged proficiency score, which was determined by L2 learning duration, age of acquisition, self-reported language exposure, and language use. We found that the influence of language proficiency on emotional prosody processing is lexicality-dependent: L2 proficiency modulated group differences for real words but not for the contrast between pseudowords and resting baseline (Figure S2) or between real words and pseudowords (Figure S3).

Within real words (Figure 5), significant group differences were found across cognitive and affective regions, including IFJa, IFJp, IFSa, IFSp, p9-46v, IP1, 55b, PEF, FOP1, and IP2, consistent with the regions identified in the native speakers and L2 learners above. Post hoc Tukey comparisons showed no significant difference between higher and lower proficiency learners in eight of these parcels (IFJa, IFJp, IFSp, p9-46v, IP1, 55b, PEF, IP2), indicating a proficiency-independent reduction in fronto-parietal engagement among L2 learners. Interestingly, in IFSa and FOP1, higher proficiency learners showed significantly stronger activation than lower proficiency learners. Given IFSa’s role in lexical-semantic processing and FOP1’s role in affective salience processing, this finding suggests that higher proficiency may be associated with greater engagement of semantic and affective mechanisms that support emotional prosody processing in real words specifically, rather than a general increase in auditory engagement.

**Figure 5.**
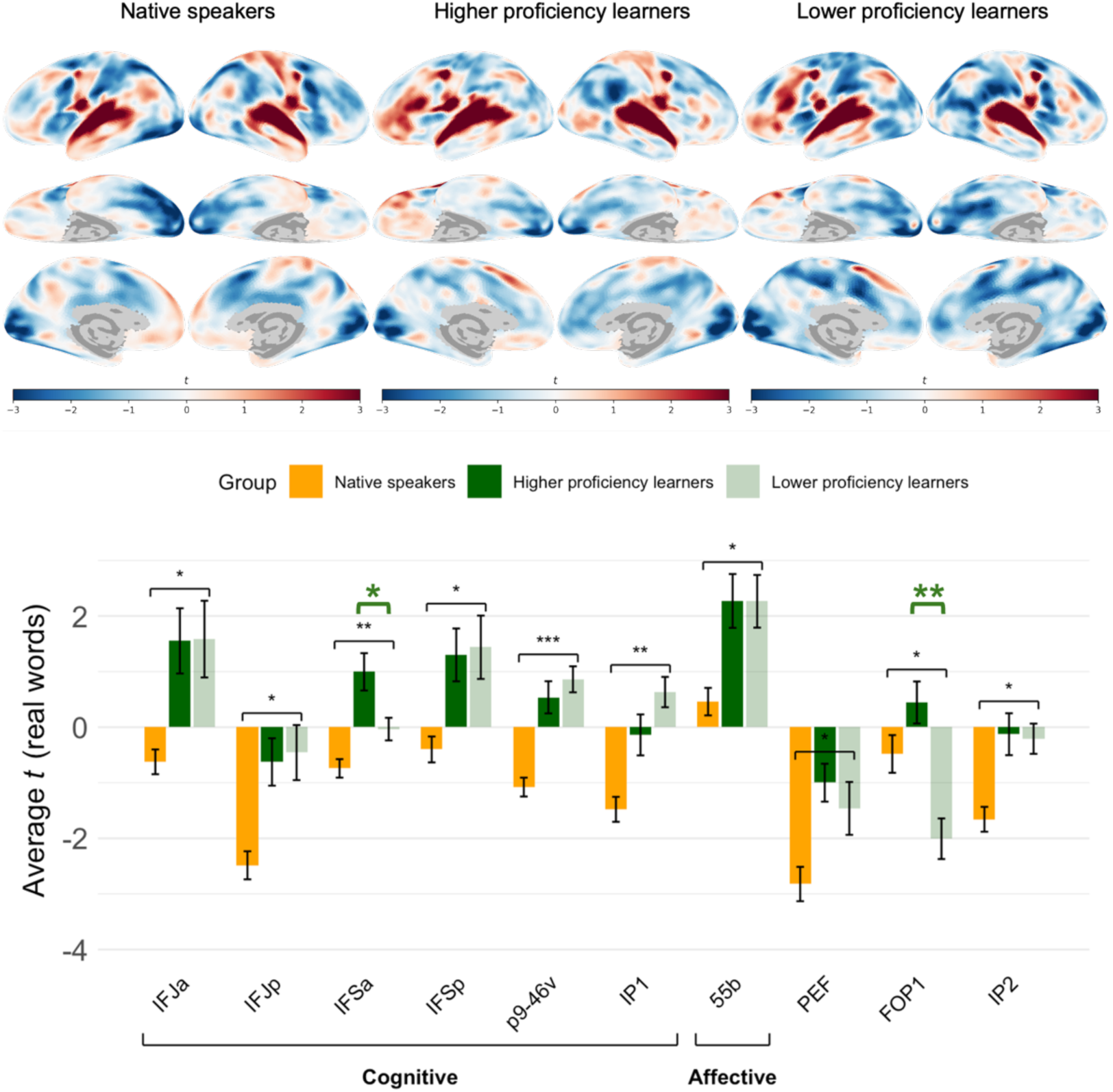
Differences between native speakers, high proficiency learners, and low proficiency learners when processing real words. (Top) Contrast maps between real words and resting baseline for native speakers, higher proficiency learners, and lower proficiency learners, where the red color indicates greater responses to real words, and blue indicates greater responses to resting baseline. (Bottom) Average contrast between real words and resting baseline in auditory, cognitive, and affective ROIs for native speakers (yellow), higher proficiency learners (dark green), and lower proficiency learners (light green). Error bars represent standard errors. Black asterisks indicate group differences between native speakers and L2 learners. Green asterisks indicate group differences between higher and lower proficiency learners. \*\*\**p* < .001, \*\**p* < .01, \**p* < .05.

## Discussion

This study investigated the neural mechanisms underlying emotional prosody processing in Mandarin Chinese by comparing native speakers and L2 learners across real-word and pseudoword stimuli. Behaviorally, while native speakers recognized emotional prosody more accurately than L2 learners, real-word stimuli were recognized faster than pseudowords in both groups. In the brain, emotional prosody processing in real words and pseudowords engaged a distributed auditory–cognitive–affective network in this tonal language. We found that group differences revealed a lexicality-dependent neural distinction. Furthermore, language proficiency modulated emotional prosody processing only for real words, with higher proficiency learners converging toward native-like activation in cognitive and affective regions in the processing of emotional prosody in real words, while no proficiency effect was observed for pseudowords. These findings provide the first neural evidence in a tonal language that lexicality is a key moderator of the in-group advantage in emotional prosody processing, helping to explain inconsistent IGA findings in the prior literature. The results are discussed below.

Consistent with Xiao and Liu ^15^, native speakers showed significantly higher accuracy than L2 learners in emotional prosody recognition in both real-word and pseudoword stimuli, replicating the IGA effect ^3,16^ in a tonal language. This IGA effect was not observed in RT. Notably, there was a main effect of lexicality, with real-word stimuli recognized faster than pseudoword stimuli, consistent with Raettig and Kotz ^43^. One possible explanation is that because these behavioral responses were collected in a block design within the scanner, participants could anticipate the likely response category before full stimulus analysis was complete, which may have attenuated the RT-level IGA while leaving accuracy intact. However, this is not the only explanation. A further possibility is that IGA is moderated by lexicality. Prior findings show a striking split by the choice of stimulus: studies using pseudoword stimuli consistently support the IGA hypothesis ^26,27,32^, whereas studies using real-word stimuli found mixed or reversed evidence patterns ^15,28,31^. Therefore, our results suggest that emotional prosody processing may draw on two partly different mechanisms for native and non-native speakers: a native speaker’s advantage in perceptual sensitivity captured by accuracy ^62,63^, and a lexically driven facilitation of response captured by RT ^64–66^.

Neuroimaging results revealed a significant interaction between group and lexicality in the processing of emotional prosody. Overall, both native speakers and L2 learners engaged in a broadly overlapping auditory, cognitive, and affective network, including the auditory cortex, inferior frontal cortex, superior temporal regions, and the temporoparietal junction. The spatial distribution and effect directions remained largely similar across positive prosody and negative prosody, suggesting a stable auditory–cognitive–affective network. This indicates that the two groups draw on shared neural architecture for processing emotional prosody. Critically, however, converging whole-brain and ROI analyses showed that the group differences within this network flipped both in direction and in anatomical locus depending on lexicality. Specifically, L2 learners showed greater frontoparietal recruitment in cognitive and affective regions for real words, whereas native speakers showed stronger temporal recruitment of auditory regions for pseudowords. This distinct pattern showed that native and L2 speakers may differ in which network, cognitive/affective or auditory, is preferentially recruited as a function of lexicality. Notably, the regions identified here (IFG, STG/STS, and premotor cortex) correspond closely to the network implicated in emotional prosody processing in prior work on non-tonal languages _1,4,7,38,39_, suggesting that the emotional prosody processing network is largely conserved across tonal and non-tonal languages. Taken together, our results demonstrate that native speakers’ advantage may be carried by a lexicality-dependent distinction between frontoparietal and temporal networks, offering a candidate mechanism for the inconsistent behavioral findings in the emotional prosody literature.

Why does lexicality modulate emotional prosody processing differently for native speakers and L2 learners? This neural distinction pattern maps closely onto established models of bilingual language processing and emotional prosody processing. Under the Revised Hierarchical Model ^23^, when processing emotional prosody in real words, L2 words lack the direct, automatized route to meaning that L1 words have, and access is instead mediated by additional lexical-conceptual computation. Integrating emotional prosody with this less automatized semantic route may therefore recruit the domain-general cognitive-control network more heavily, where prefrontal and parietal regions are repeatedly implicated in bilingual language control ^67^. This is consistent with the stronger activation we observed in L2 learners relative to native speakers in the inferior frontal cortex, dorsolateral prefrontal cortex, the inferior parietal and temporo-parieto-occipital junction regions, and the music perception hub for real words. Pseudowords, by contrast, offer no lexical-semantic route: because emotional prosody must be extracted from the acoustic signal, the group difference may thus reduce to a difference in auditory analysis of native versus non-native prosodic patterns, localized to auditory regions including primary/belt auditory cortex (A4, A5, TA2) and superior temporal regions (STGa, STSdp). These auditory regions align well with the three-stage sequential model of emotional prosody proposed by Schirmer and Kotz ^4^ and the dual-stream model of speech processing ^68,69^.

Notably, the stronger involvement of auditory regions is not purely acoustic. Because pseudoword stimuli used in the current study were constructed to be phonotactically legal and word-like, they likely partially activated a set of real phonological neighbors in the listener’s lexicon, which must then be resolved or ruled out before the item is rejected as a non-word ^66,70^. Native speakers, with a large and dense native lexicon, would activate more neighboring words in response to a word-like pseudoword, engaging temporal-lobe regions more heavily in this competition resolution process. However, L2 learners, with a smaller and sparser L2 lexicon, would have fewer neighbors to activate and resolve, yielding weaker engagement of the same region. This is consistent with recent evidence that STG jointly encodes acoustic-phonetic and lexical information in constructing word-form representations ^71^. A parallel pattern emerged for real words: native speakers showed stronger inferior parietal lobule activation than L2 learners, a region implicated in semantic association and integration. Taken together, these results suggest that the native speakers’ emotional prosody processing advantage in pseudowords may also reflect a richer and more efficient engagement of lexico-semantics, rather than a difference in low-level auditory perception alone; L2 learners’ comparatively limited lexical knowledge constrains their capacity to engage these integrative and competition–resolution mechanisms.

Finally, second language proficiency modulated emotional prosody processing selectively for real words, not for pseudowords. Among L2 learners, greater proficiency was associated with stronger engagement of IFSa and FOP1 during emotional prosody processing in real words, whereas this proficiency effect was absent for pseudowords. This pattern is consistent with the single-system accounts of lexical processing ^72,73^, in which cumulative exposure gradually strengthens lexical representations within a unified system, such that proficiency effects emerge specifically where lexical representations are sufficiently consolidated to support active retrieval. Real words have well-established, item-specific lexical representations, including phonological form and meaning, allowing proficiency to modulate the degree of lexico-semantic engagement. Pseudowords, by contrast, have no such stored representation regardless of proficiency levels: because both groups encountered these unfamiliar phonological forms for the first time, there is no accumulated lexical knowledge for proficiency to differentiate, and thus the two learner groups pattern together rather than diverging. This finding suggests that what is acquired with increasing L2 experience is not a generic improvement in auditory sensitivity, but specifically the lexico-semantic scaffolding that supports emotional prosody processing when lexical content is available. Given the limited sample size of the current study, further investigation is needed to determine how proficiency shapes these neural patterns.

To conclude, our findings show that lexicality shapes native and non-native speakers in fundamentally different ways. For native speakers, emotional prosody processing draws flexibly on both lexico-semantic integration and acoustic–perceptual analysis. For non-native speakers, it depends more heavily on effortful, domain-general cognitive control. Furthermore, language proficiency modulated emotional prosody processing only for real words, but not for pseudowords. Together, lexicality is not a peripheral stimulus property but an important factor in bilingual emotion processing, shaping both how native and non-native speakers diverge and how the degree of that divergence varies with L2 experience.

## Methods

## Participants

Thirty right-handed, neurologically healthy young adults from the University of South Carolina community participated in the study, including 15 native Chinese speakers (Native Group, 9 females, *M*_age_ = 24.4 years, age range = 20–28) and 15 L1-English L2-Chinese learners with at least six months of Chinese learning experience (Learner Group, 9 females, *M*_age_ = 21.3 years, age range = 18–32). All 15 participants in the Learner Group were native English speakers who were enrolled in a Chinese class and had completed at least one semester of Chinese study. They all scored 80% or higher on a vocabulary screening test of the real-word stimuli administered within one week prior to the fMRI experiment (Mean ACC = 91.9%). Neither group had any prior knowledge of the pseudowords, ensuring that both groups had similar semantic knowledge of the real words and pseudowords. None of the L2 Chinese learners were heritage speakers of Mandarin Chinese or any other tonal language, All participants had normal or corrected-to-normal vision and reported no hearing problems. All participants provided informed consent and received moderate monetary compensation. The study procedures were approved by the Institutional Review Board (IRB) of the University of South Carolina.

## Stimuli and design

The stimulus set included three semantic valence categories: 48 pseudowords with neutral semantic valence, 48 real words with negative semantic valence, and 48 real words with positive semantic valence. All words were matched for syllable length and lexical tone, and positive and negative words were further matched for word frequency ^74^, valence ^75^, arousal ^75^, concreteness _76_, and age of acquisition ^77^. Pseudoword stimuli were constructed as disyllabic lexical gaps ^78^, where each syllable is phonotactically legal yet does not correspond to any existing lexical entry in Mandarin Chinese. For example, “chun” can occur with tone 1, 2, and 3, but not tone 4, making “chun^4^” a lexical gap in Chinese; each pseudoword combined two such syllables. Furthermore, we created the sentence stimuli by embedding each word stimulus in the carrier sentence “ta^1^ hen^3^ (*She is very*),” resulting in 48 pseudoword sentences with neutral semantic valence, 48 real-word sentences with negative semantic valence, and 48 real-word sentences with positive semantic valence. The carrier sentence was chosen to minimize contextual and syntactic processing ^48^. Table 1 presents examples of real-word and pseudoword stimuli with negative, positive, and neutral semantic valence. Table 2 presents the means and standard deviations of relevant psycholinguistic variables in positive and negative word sets.

**Table 1.** Examples of real-word and pseudoword stimuli for Chinese words and sentences.

|  | **Real Word** |  | **Pseudowords** |
| --- | --- | --- | --- |
| Semantic valence | Negative | Positive | Neutral |
| Word | ao<sup>4</sup>man<sup>4</sup><br>arrogant<br>“arrogant” | rui<sup>4</sup>zhi<sup>4</sup><br>wise<br>“wise” | chun<sup>4</sup>pou<sup>4</sup><br>chunpou<br>“chunpou” |
| Sentence | ta<sup>1</sup> hen<sup>3</sup> ao<sup>4</sup>man<sup>4</sup><br>She very arrogant<br>“She is very arrogant.” | ta<sup>1</sup> hen<sup>3</sup> rui<sup>4</sup>zhi<sup>4</sup><br>She very wise<br>“She is very wise.” | ta<sup>1</sup> hen<sup>3</sup> chun<sup>4</sup>pou<sup>4</sup><br>She very chunpou<br>“She is very chunpou.” |

**Table 2.** Means (standard deviations) of psycholinguistic variables in negative and positive words. All differences were n.s., all p > 0.1 (Welch’s two-sample *t*-test).

|  | **Negative words** | **Positive words** |
| --- | --- | --- |
| Frequency | 17.73 (29.85) | 17.25(25.61) |
| Valence | -1.78 (0.41) | 1.78 (0.38) |
| Arousal | 2.50 (0.51) | 2.44 (0.43) |
| Concreteness | 3.36 (0.31) | 3.36 (0.31) |
| Age of Acquisition | 11.49 (1.81) | 10.91 (1.74) |

Finally, we created multiple audio clips for each word or its corresponding sentence stimulus. Two native Chinese speakers, one female and one male, recorded all stimuli. The same word was recorded twice by each speaker, once with positive emotional prosodic valence (e.g., joy), and once with negative emotional prosodic valence (e.g., sad). All the stimuli were recorded in a soundproof studio using Audacity (Version 3.0.0). The stimuli were segmented and normalized to the same amplitude using Praat (Version 6.3.10).

In summary, the auditory stimuli comprised a total of 1152 items: 192 pseudowords and pseudoword sentences with negative prosody and neutral semantic valence, 192 pseudowords and pseudoword sentences with positive prosody and neutral semantic valence, 192 real words and real-word sentences with negative prosody and positive semantic valence, 192 positive real words and real-word sentences with positive prosody and positive semantic valence, 192 real words and real-word sentences with negative prosody and negative semantic valence, and 192 real words and real-word sentences with positive prosody and negative semantic valence. The emotional prosody of all selected and normalized recordings was validated by 42 native Chinese speakers (23 females, 19 males) in an emotional prosody rating task. The rating results confirmed that each stimulus’s emotional prosody had its expected prosodic valence ratings and that the two sets (i.e., positive and negative) of emotional prosody stimuli had comparable arousal ratings (Figure S1).

## Imaging procedure and sequence parameters

MRI data were acquired using a Siemens 3T Prisma Fit scanner equipped with a 20-channel head coil at the McCausland Center for Brain Imaging at the University of South Carolina. Participants lay on their backs and wore a glove response box on their right hand, with their legs supported by a wedge pillow and a blanket provided for warmth if requested. Foam cushions were used to stabilize the position of the head inside the head coil and to minimize motion. During scanning, auditory stimuli were presented via E-Prime 3.0 software (Psychology Software Tools, Pittsburgh, PA) through Optoacoustics MRI-compatible headphones. The sound level was adjusted to the highest level comfortable for each participant. Figure 6 illustrates the experiment design and MRI protocol.

**Figure 6.**
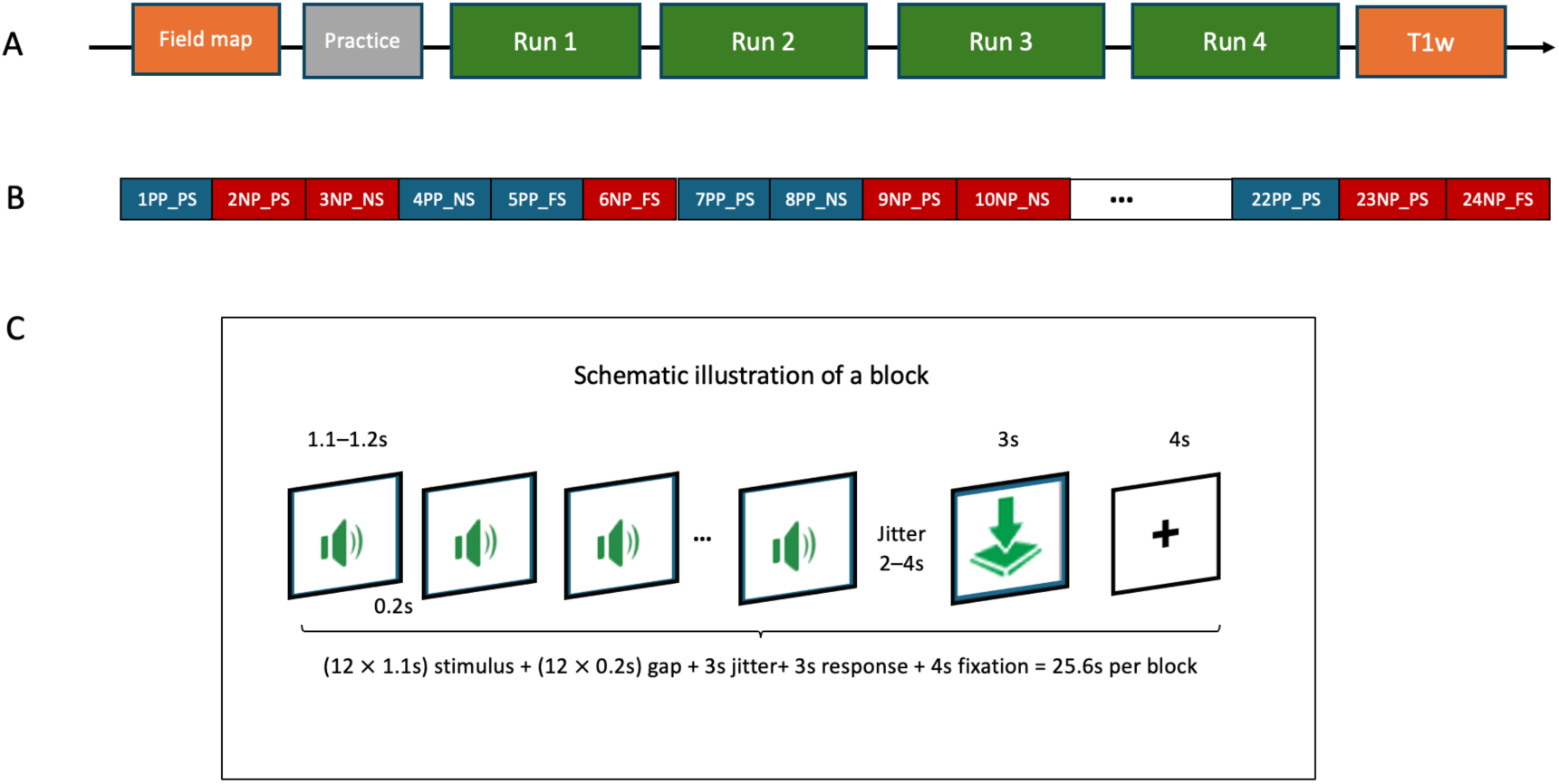
Illustrations of the MRI scanning flowchart (A), an example of a functional run (B) with 24 blocks (red: negative prosody; blue: positive prosody), and the schematic illustration of a block (C). The order of blocks was randomized within the run, and the order of runs was counterbalanced across participants. PP: positive emotional prosody; NP: negative emotional prosody; PS: positive semantics; NS: negative semantics; FS: neutral semantics.

The main experiment included four functional runs, corresponding to the combinations of speaker gender (male, female) and stimulus length (word, sentence). The order of the four runs was counterbalanced across participants (Figure 6A). Within each run, a block design with six conditions was used, consisting of twenty-four randomized blocks (Figure 6B). Each block lasted about 26 seconds. In each block (Figure 6C), participants listened to twelve randomized auditory stimuli (each 1.1–1.2 s), separated by a 200ms silence gap to prevent overlap. After the twelve stimuli, a jittered fixation period (2–4 s) was presented, followed by a 3-second response phase during which participants used an MRI-compatible response glove to indicate whether the speaker sounded like they were experiencing negative or positive emotions. The mapping of the finger presses for positive or negative responses was counterbalanced across participants. A 4-second fixation followed the response phase before the next block began. Before the fMRI scans, two field maps and a short practice run were acquired to familiarize participants with the task and response device; at the end of the session, a high-resolution T1-weighted anatomical scan was obtained (Figure 6A).

Functional images were acquired using a T2*-weighted multiband echo-planar imaging (EPI) sequence with the following parameters: field of view (FOV) = 215 × 215 × 135 mm^3^, voxel size = 3 × 3 × 3 mm^3^, repetition time (TR) = 1500 ms, echo time (TE) = 26.2 ms, slices = 45, flip angle (FA) = 44°. Each word run included 440 volumes and lasted approximately 11 minutes, and each sentence run included 454 volumes and lasted approximately 11.5 minutes. The T1-weighted anatomical scan was acquired using a multi-echo MPRAGE sequence: FOV = 256 × 256 × 192 mm^3^, voxel size = 1 × 1 × 1 mm^3^, TR = 2530 ms, TEs = 1.44, 2.9, 4.36, 5.82, 7.28 ms, FA = 7°. The T1w scan provided whole-brain coverage and took 6 minutes to complete. The total scanning time was approximately 1 hour.

## Analysis

Neuroimaging data were preprocessed using fMRIPrep ^55^ version 25.1.4 and resampled onto the onavg-ico32 cortical surface template ^79^. We further reduced noise by regressing out nuisance regressors from the resampled data and by z-scoring the residual time series. Similar to Feilong, et al. ^80^, nuisance regressors were partialed out from functional data separately for each run. These regressors included six motion parameters and their derivatives, framewise displacement _81_, six principal components from cerebrospinal fluid and white matter (aCompCor) ^82^, and polynomial trends up to second order. The residual time series of each surface vertex in each run was normalized to zero mean and unit variance.

The same brain response pattern may have different topographies on different brains ^57^, which limits the power of statistical comparisons. To resolve these topographic idiosyncrasies across individuals, we functionally aligned all participants using hyperalignment ^56,57^. In particular, we used connectivity hyperalignment ^83^ because it does not require the stimulus presentation to be time-locked across participants. Specifically, for each of the 30 participants, we used two runs to derive individual hyperalignment transformations to a common representational space and applied these transformations to the remaining two runs for statistical analysis. The two test runs were selected per participant as the pair with the highest inter-run correlation across all possible run pairings. Because word runs and sentence runs did not differ significantly, they were treated as equivalent for this purpose (see details in Supplemental Methods). All analyses reported in the Results are based on these two selected test runs. Data were then smoothed using a 10-mm FWHM kernel prior to GLM estimation.

For each block, accuracy (ACC) was binary coded: 1 for correct judgments of emotional prosody and 0 for incorrect judgments. For reaction time (RT), we excluded trials with incorrect judgments, RTs shorter than 200 ms, and RTs that exceeded three standard deviations from each participant’s mean. Logistic and linear mixed-effects models were used to analyze ACC and RT, respectively, with participant included as a random intercept ^84^ using R (R Core Team, 2024). The full models for ACC and RT analyses included two fixed factors: (1) group with two levels (i.e., native group and learner group), (2) lexicality with two levels (i.e., real-word stimuli and pseudoword stimuli), dummy-coded. We used a maximal mixed-effect model that considered all potential main effects and interactions ^85^. Statistical analyses were performed in R (R Core Team, 2024). In addition, behavioral and neuroimaging data were analyzed in Python using the neuroboros package.

We conducted region-of-interest analyses across all 180 cortical parcels of the HCP-MMP1.0 atlas ^86^. For each parcel, we extracted the mean activation *t*-value per participant across the vertices for several contrasts of interest (e.g., real word vs. pseudoword, real word vs. rest, pseudoword vs. rest), separately for positive and negative prosody where applicable. Independent-samples *t*-tests compared each contrast between the Native and Learner groups at every parcel, and the resulting p-values were corrected for multiple comparisons across the 180 parcels using the false discovery rate (FDR) ^87^. Parcels surviving FDR correction at *p* < .05 were considered significant, and those at *p* < .1 were considered marginally significant. Parcels were labeled according to their corresponding HCP-MMP1.0 region name for interpretation.

To further investigate whether the neural response to lexicality varies as a function of language proficiency, we divided L2 learners into two groups (i.e., higher vs. lower proficiency learners) based on their averaged proficiency score, which was determined by L2 learning duration, age of acquisition, self-reported language exposure, and language use. Whole-brain and ROI analyses were also conducted across three groups: native group, higher proficiency learners, and lower proficiency learners. For each parcel, each participant’s mean t-value was computed by averaging across all vertices. A one-way ANOVA (Native vs. L2 high vs. L2 low) was conducted independently for each parcel, and the resulting p-values were corrected for multiple comparisons across parcels using the Benjamini-Hochberg FDR procedure (q < .05). For any parcel showing a significant omnibus group effect after FDR correction, pairwise post hoc comparisons among the three groups were conducted using Tukey’s Honestly Significant Difference (HSD) test to identify which specific group pairs differed.

