## Supplemental Methods, Figure S1, Figure S2, Figure S3 for "Brain Mechanisms of Lexicality in Bilingual Emotion Processing"

***Supplemental Materials* for Brain Mechanisms of Lexicality in Bilingual Emotion Processing**

### Supplemental Methods

Each participant completed four functional runs: two word runs and two sentence runs. For each participant, we compared within-task similarity to between-task similarity separately, based on the contrast between real-word and pseudoword conditions. Within-task similarity was defined as the average of the Fisher-z-transformed correlation between the two word runs and between the two sentence runs. Between-task similarity was defined as the average Fisher-z-transformed correlation across all word-to-sentence-run pairings. Paired-sample t-tests showed no difference between within-task and between-task similarity across subjects ($t = -0.928$, $p = .361$), indicating no evidence of task-specific structure in the inter-run correlations. To assess reliability across groups, we compared the inter-run correlation of the two selected runs between native speakers and L2 learners and found no significant group difference ($t = -0.501$, $p = .620$).

### Supplemental Figures

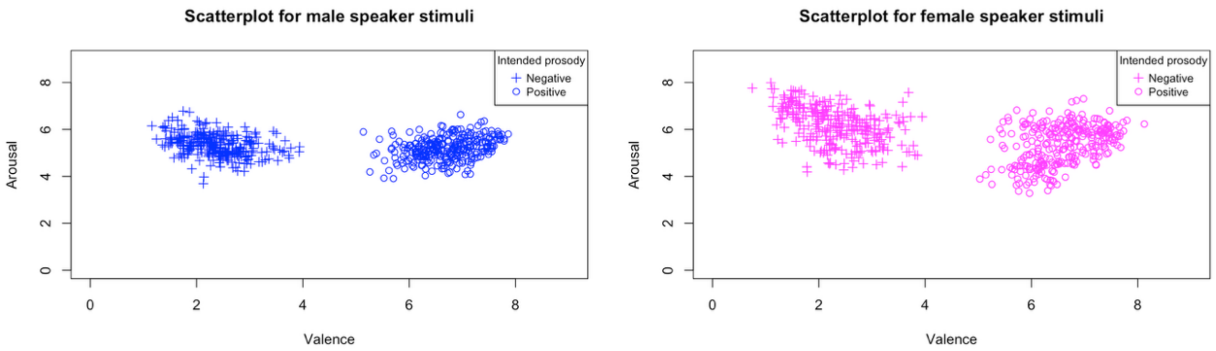


**Figure S1.** Scatterplots of the mean valence and mean arousal ratings for each positive and negative stimulus produced by male (blue) and female (magenta) speakers on a scale from 0 to 9. On the X-axis (valence), 0 indicates "very negative," and 9 indicates "very positive." On the Y-axis (arousal), 0 indicates "very excited," and 9 indicates "not excited." A cross (+) represents that the stimulus was intended to convey negative emotional prosody, and a circle (O) represents intended positive emotional prosody. All stimuli with negative prosody had ratings below 4 (valence: $M = 2.34$, $SD = 0.61$), and all stimuli with positive prosody had valence ratings above 5 (valence: $M = 6.65$, $SD = 0.58$). No stimulus had a valence rating between 4 and 5, which would indicate neutral prosodic valence. The positive and negative prosody types were matched in arousal ratings (negative prosody: $M = 5.76$, $SD = 0.79$; positive prosody: $M = 5.27$, $SD = 0.69$).

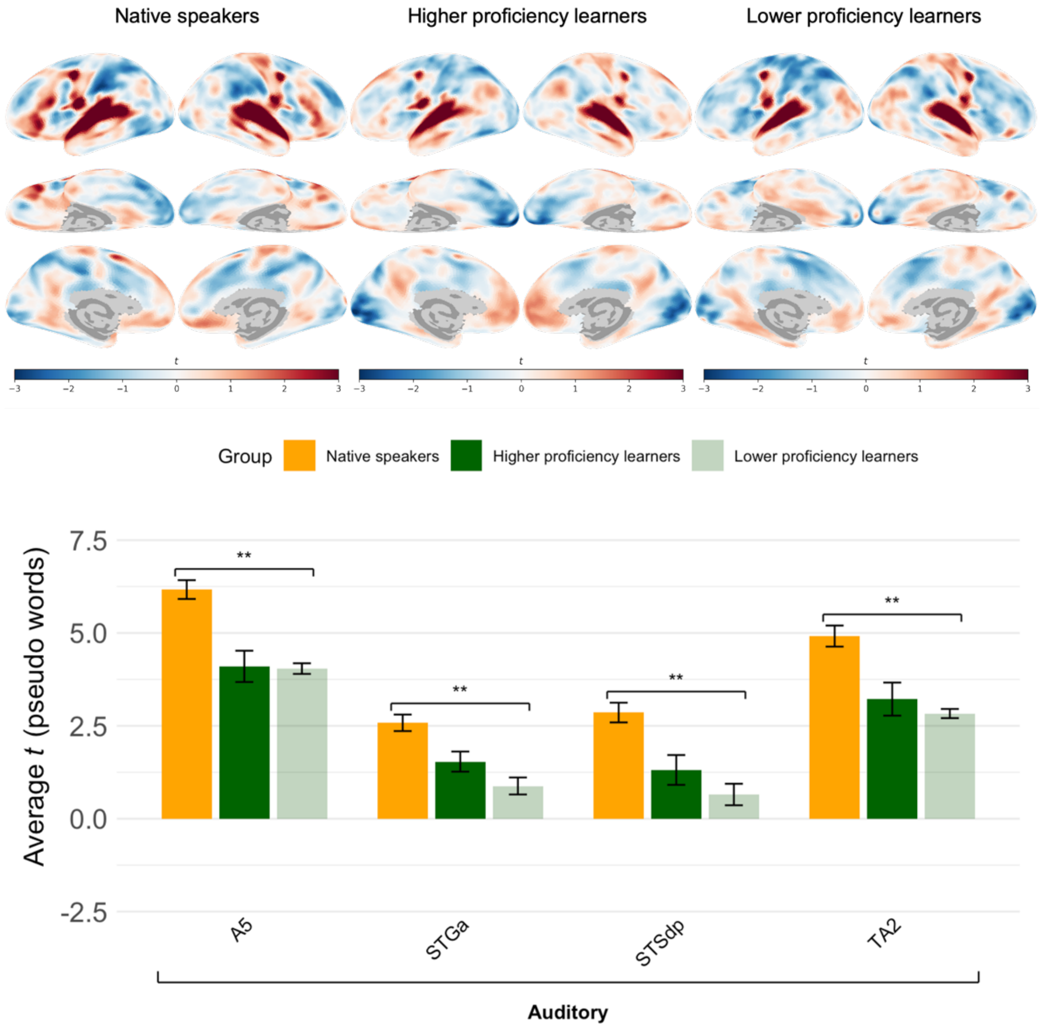


**Figure S2.** Differences between native speakers, higher proficiency learners, and lower proficiency learners when processing pseudowords. (Top) Contrast maps between pseudowords and resting baseline for native speakers, higher proficiency learners, and lower proficiency learners, where the red color indicates greater responses to pseudowords, and blue indicates greater responses to the resting baseline. (Bottom) Average contrast between pseudowords and rest in auditory, cognitive, and affective ROIs for native speakers (yellow), higher proficiency learners (dark green), and lower proficiency learners (light green). Error bars represent standard errors. Black asterisks indicate group differences between native speakers and L2 learners. No group difference was found between higher and lower proficiency learners. ***$p < .001$, **$p < .01$, *$p < .05$.

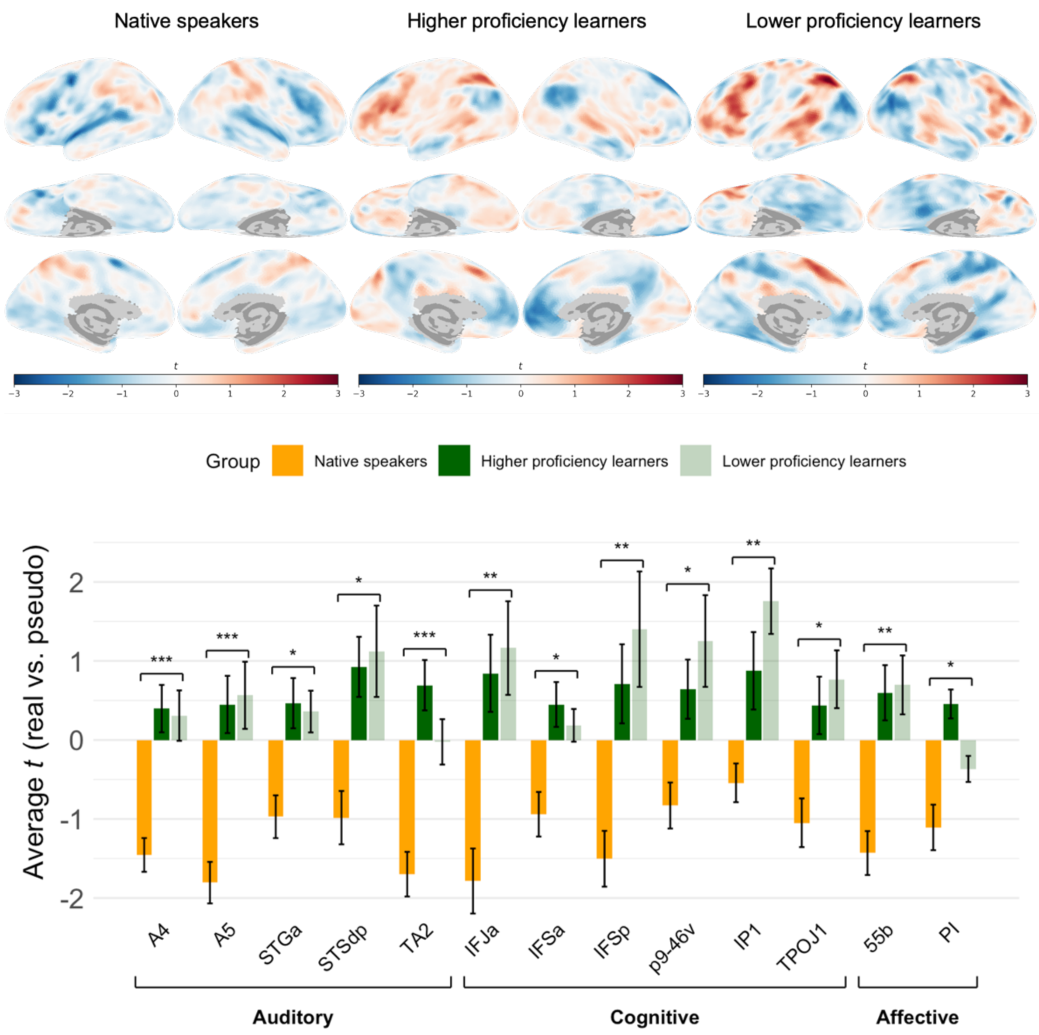


**Figure S3.** Differences in the real-word and pseudoword contrast across native speakers, higher proficiency learners, and lower proficiency learners. (Top) Contrast maps between real words and pseudowords for native speakers, higher proficiency learners, and lower proficiency learners, where the red color indicates greater responses to real words, and blue indicates greater responses to pseudowords. (Bottom) Average contrast between real words and pseudowords in auditory, cognitive, and affective ROIs for native speakers (yellow), higher proficiency learners (dark green), and lower proficiency learners (light green). Error bars represent standard errors. Black asterisks indicate group differences between native speakers and L2 learners. No group difference was found between higher and lower proficiency learners. ***$p < .001$, **$p < .01$, *$p < .05$.
